# Chronic *Toxoplasma gondii* infection curtails the cytotoxic potential of acute T cell responses to West Nile virus in the brain

**DOI:** 10.1101/2024.01.21.576564

**Authors:** Jennifer L. Uhrlaub, Kathryn E. McGovern, Emily F. Merritt, Michael S. Kuhns, Anita A. Koshy, Janko Ž. Nikolich

## Abstract

*Toxoplasma gondii (T. gondii)*, a common brain-tropic parasite, chronically infects the central nervous system (CNS) of up to a third of the world’s population. Constant immune surveillance interrupts cyst reactivation within the CNS and dramatically alters the immune landscape of the brain. West Nile virus (WNV) is a mosquito-borne infection with a clinical spectrum ranging from asymptomatic to mild flu-like symptoms to severe neuroinvasive disease. In a cohort of WNV infected people, we discovered a positive correlation between WNV disease severity and *T. gondii* seropositivity. In a mouse model pairing chronic *T. gondii* with acute WNV infection, we found an increased susceptibility of mice to WNV, with reduced granzyme B expression in WNV-specific T cells and increased regulatory T cell (Treg) numbers in the brain, but not the periphery. This demonstrates that the *T. gondii*-infected tissue microenvironment impairs immune defense against other brain infections by blunting local T cell responses.

## Introduction

*Toxoplasma gondii* (*T. gondii*) is an obligate intracellular parasite that naturally infects a wide range of warm-blooded hosts including humans and rodents [1]. It is one of the most prevalent parasites globally with seropositivity in some regions of the world above 60% [1–3]. People are commonly infected with *T. gondii* through the consumption of contaminated food and water. In the United States, *T. gondii* is considered a neglected parasitic infection and is a leading cause of death from foodborne illness [4]. Approximately 40 million individuals, or 11% of the total U.S. population older than 6 years of age, have been infected with *T. gondii* [4].

During acute infection, *T. gondii* rapidly proliferates and disseminates throughout the body as a tachyzoite. T cell-mediated defenses limit tachyzoite expansion primarily through the production of IFNγ [5–7]. In select organs and cells, a small population of parasites differentiate into slower-growing bradyzoites, which form intracellular cysts that can endure for the life of the host. The central nervous system (CNS) is a major organ in which parasites encyst and persist, but the cost of *T. gondii* carriage to the host remains incompletely understood. *T. gondii* quiescence is maintained by a dynamic and local immune response dominated by high levels of IFNγ produced by tissue-resident CD8 T cells [8]. Cytolytic activity is a dispensable mechanism for parasite control during acute toxoplasmosis but may play a role during chronic infection to prevent recrudescence and clear infected neurons [9,10]. This is a delicate balance, and the parasite has evolved its own immune evasion mechanisms that limit host cell death, including granzyme mediated apoptosis[11]. While these *in vitro* studies by Yamada et al. examined *T. gondii* inhibition of granzyme function within infected cells [11], whether *T. gondii* infection may also limit the differentiation and polarization of T cells has not been investigated.

West Nile virus (WNV) is a flavivirus transmitted primarily by mosquitoes. The reemergence of WNV in North America in 1999 coincided with its proclivity to cause severe neurological disease [12]. After initial infection via the mosquito bite, the virus enters the bloodstream and eventually the brain, where it can cause inflammation and damage to neurons, leading to a range of neurological symptoms. Most WNV infections are asymptomatic, with∼15% causing fever and ∼5% resulting in severe and life-threatening encephalitic disease, dominantly in older adults and those with underlying health conditions or weakened immune systems[13]. Because *T. gondii* can establish a lifelong infection in the CNS and WNV is also a neurotropic infection, these two microbial pathogens would be expected to occasionally coinfect humans.

T cells play a crucial role in WNV clearance. In mouse models, these cells arrive in the brain on days 8-9 post-infection and control viral load by producing antiviral cytokines or by triggering death of infected cells through perforin- or Fas ligand-dependent pathways [14,15]. Perforin and granzyme are two key molecules involved in the cytolytic activity of CD8+ T cells and are required to clear WNV from infected neurons [14]. The expression of perforin and granzymes positively correlates with reduction in viral RNA levels, supporting that cytotoxic CD8+ T cells clear WNV infection in the brain [16]. Th1 CD4 T cells also contribute to the clearance of WNV via cytolytic mechanisms [17]. Overall, the cytolytic activity of T cells, particularly through the use of perforins and granzymes, is an important mechanism to control WNV infection. To investigate the impact of *T. gondii* persistence, and tissue resident memory, on the *de novo* T cell response to acute WNV, we have developed a *T*.*gondii*/WNV co-infection mouse model. Our data reveal that *T. gondii* alters the CNS environment, reducing the differentiation of T cells expressing granzyme B (GzmB), and decreasing survival from WNV infection. Some of that effect can be ascribed to increased numbers of resident regulatory T cells (Treg) in the brain during chronic *T. gondii* persistence. These data inform our understanding of immune responses within a brain colonized by a parasite and provide evidence that the dynamics of immune defense in the brain is shaped by tissue infectious history of the host.

## Results

Given that immune responses in the brain must both protect the host and largely spare normal function of the organ, we hypothesized that mechanisms which prevent pathology and disease during *T. gondii* persistence may limit immune responses within the CNS, potentially worsening outcome following infection with a neurotropic virus. To test this hypothesis, we first explored potential clinical association between WNV disease severity and *T. gondii* infection, by determining seroprevalence of *T. gondii* antibodies in a cohort of individuals who experienced WNV infection. We found that more than half of individuals diagnosed with West Nile encephalitis were co-infected with *T. gondii*, which was a significantly higher prevalence than in those with West Nile fever (*p*=0.0047, **Table I;** a handful of participants in the fever cohort were also diagnosed with viral meningitis, a generally benign disease). While these results originate from a limited number of samples, they suggest that *T. gondii* persistence may be a risk factor for increased neurological disease following an infection like West Nile virus.

**Table I.** *T. gondii* seropositivity is associated with worse neurological symptoms from WNV. Seropositivity for *T. gondii* was measured by ELISA as described in *Methods*. Participants were clinically defined for WNV disease severity ranging from West Nile fever (WNF), West Nile encephalitis (WNE), with some in each category also having meningitis (+/- M). Fisher exact test comparing WNF to WNE, *p*=0.0043.

|  | WNF +/- M | WNE +/- M |
| --- | --- | --- |
| Total | n=37 | n=41 |
| *T. gondii* seropositive | 7 | 21 |
| % seroprevalence | 19% | 51% |
| Age (years ±SEM) | 57 ± 14 | 69 ± 12 |
| sex | n=15 male<br>n=22 female<br>n=1 unknown | n=31 male<br>n=9 female |
| race | n=29 white | n=37 white |

To address the causality between *T. gondii* infection and WNV susceptibility, we infected C57BL/6 mice with *T. gondii* and monitored survival for 60 days; mice were then challenged with WNV and monitored for an additional 40 days (**Fig. 1**). Survival of *T. gondii* infection was 48% at day 60 and WNV infection further reduced overall survival to 15%. By contrast, mice only infected with WNV exhibited a survival rate of 63% (**Fig. 1**). Therefore, our data demonstrates the synergistic and negative impact of co-infection on mouse survival.

**Figure 1.**
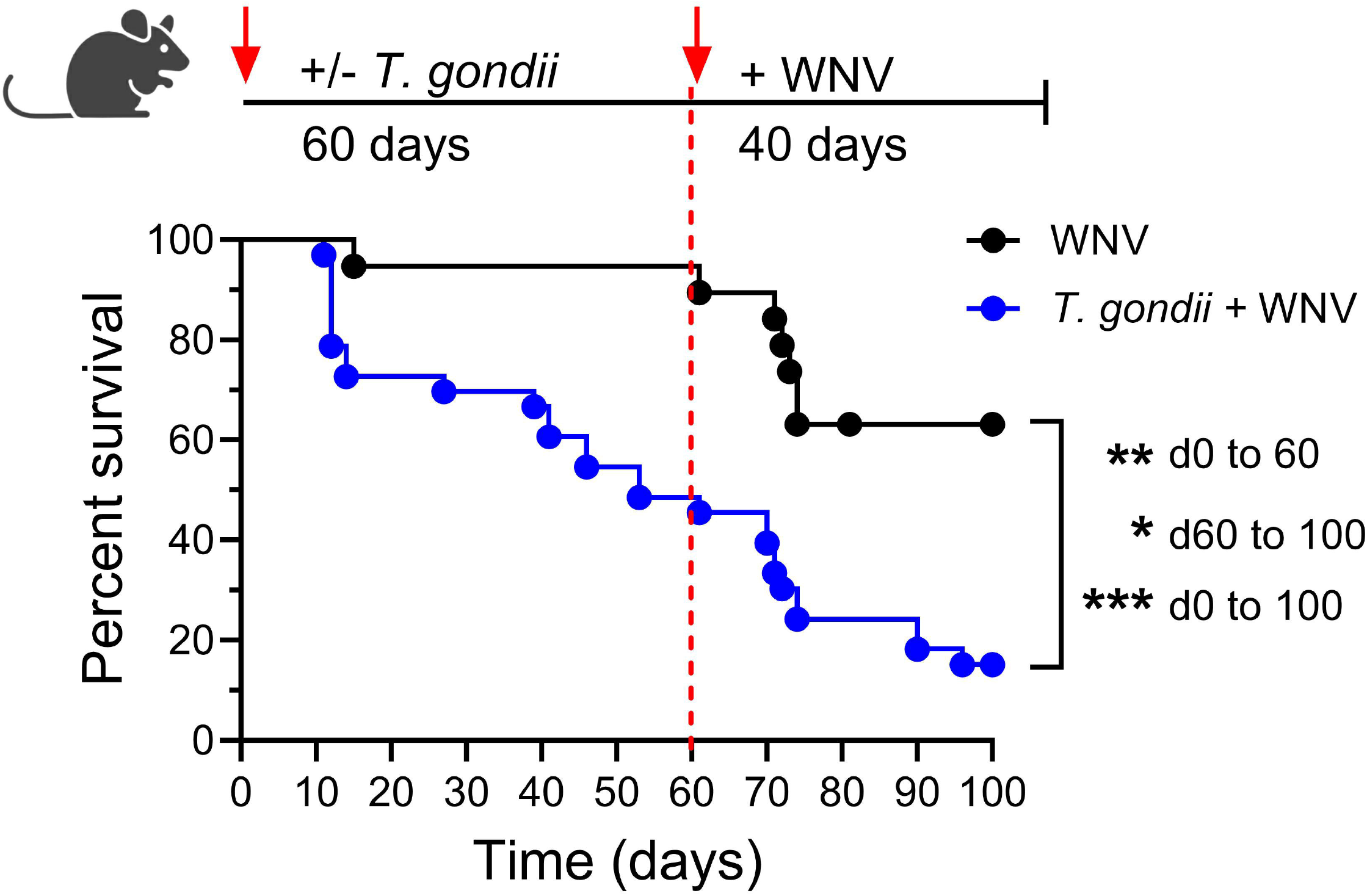

We next determined whether there was a direct effect of persistent *T. gondii* infection on viral burden and immune response to WNV over both short- and long-terms. To that effect, we interrogated a cohort of mice infected with *T. gondii* for 60 days as in **Fig. 1** and compared it to a cohort that had survived *T. gondii* infection for more than 500 days (lifelong cohort; **Supp. Fig. 1**). In a similar design, McGovern et al. showed that C57BL/6 mice eventually clear *T. gondii* as they found no evidence of parasite DNA at 600 days post-infection [18]. In our lifelong cohort, PCR analysis showed that most, but not all (9/14 mice) cleared *T. gondii* DNA in the brain.

Since some of our day 500 survivors did not clear the parasite, future studies to determine the contribution of parasite presence will be necessary.

We challenged both cohorts of mice with WNV as depicted in **Fig. 2A**. Mice were sacrificed on day 10 post-WNV infection and assessed for viral burden and cellular immune responses in the brain. We hypothesized that higher WNV titers in the brain of co-infected mice would correlate with increased susceptibility of the *T. gondii* infected mice to WNV. As previously reported [19], we did measure a significantly higher viral load in lifelong infected, older, animals (**Fig. 2B**).

**Figure 2.**
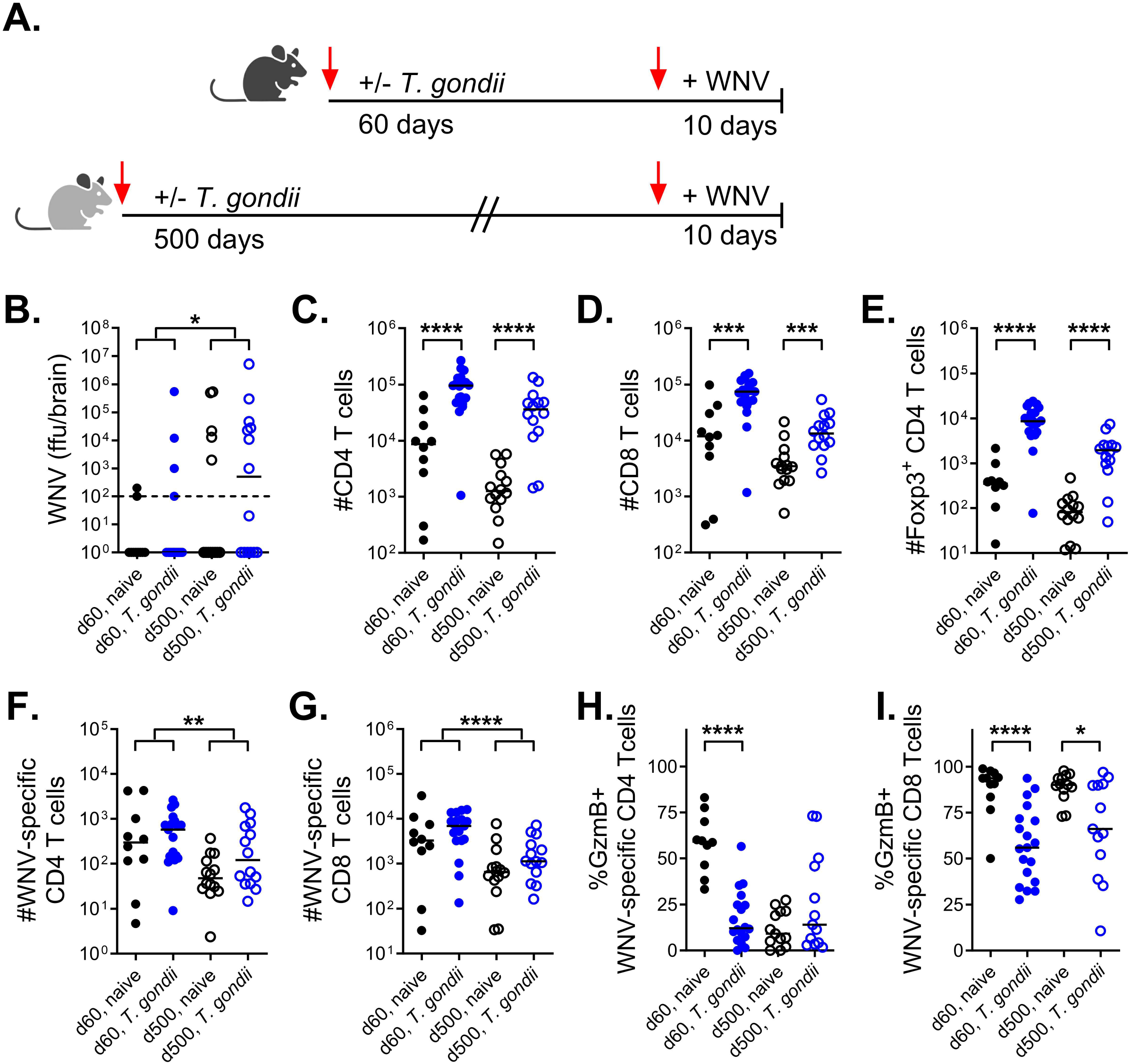

However, we determined that neither the short-term nor lifelong cohorts exhibited higher WNV titers in the brain as compared to their *T. gondii-*naïve controls (**Fig. 2B**). We conclude that increased susceptibility of co-infected mice to WNV is not directly linked to viral burden.

This finding allowed us to accurately compare how the local tissue resident immune response to *T. gondii* impacts T cells responding to WNV. Indeed, disproportionate viral burdens would confound comparisons of tissue-resident cellular immune responses as antigen load would then be the most likely driving factor for discrepancies between groups.

*T. gondii* persistence is controlled within the brain by a dynamic immune response. Local IFN-γ production is known to be essential to maintain parasite quiescence [8,20]. The introduction of a new, neurotropic, viral infection has the potential to tip this balance in multiple ways. We focused our analysis on T cells, assessing the impact of *T. gondii* on the WNV-specific immune response. We measured this via flow cytometric (FCM) analysis of T cells in the brain and compared them to those in superficial cervical lymph nodes, as a proxy for peripheral responses, on day 10 post-WNV infection. We found that in both the d60 and d500 cohorts *T. gondii* infection resulted in an increased total number of CD4 and CD8 T cells in the brain (**Fig. 2C, D**). We further found a significantly higher number of Foxp3^+^ CD4 T regulatory cells (Tregs) in the brain in the presence of *T. gondii* infection (**Fig. 2E**). The higher numbers of CD4, CD8, and CD4 Foxp3+ Tregs in the brain of *T. gondii* infected mice reflect the tissue resident memory populations established during *T. gondii* infection (**Fig. 2C, D, E**). These results are consistent with long-standing literature demonstrating that T cells are required to maintain *T. gondii* in a quiescent state [8,21]. To enumerate the antigen-specific CD4 and CD8 T cell response to WNV, we used MHC:peptide tetramers containing the immunodominant T cell epitopes, I-A^b^:ENV_641_ and K^d^:NS4b_2488_ (**Fig. 2F, G**). The numbers of these WNV-specific T cells were comparable between *T. gondii*-infected and naïve groups within each cohort, indicating that *T. gondii* persistence, and the ongoing or tissue resident immune response to the parasite does not prevent or limit the entry of WNV-specific T cells to the brain. However, we found that significantly fewer WNV-specific T cells in the brain express GzmB, a key effector of cytolytic activity, in the *T. gondii*-infected groups (**Fig. 2H, I**). This suggests that *T. gondii* must directly (immune evasion/parasite factors) or indirectly (by altering brain microenvironment) impact the differentiation of *de novo* T cell responses.

To determine whether WNV-specific T cells continue to acquire their effector function within the brain, we examined the same parameters in peripheral lymph node (LN) T cells.

Pairing LN and brain data revealed that the percentage of WNV-specific CD8 or CD4 T cells expressing GzmB continues to increase after entry to the brain (**Fig. 3A, B**). This was statistically significant for WNV-specific CD8 and CD4 T cells across all groups for both cohorts (**Fig. 3**).This finding is consistent with data showing that highly oligoclonal WNV -specific T cells continue to differentiate in the tissue [16].

**Figure 3.**
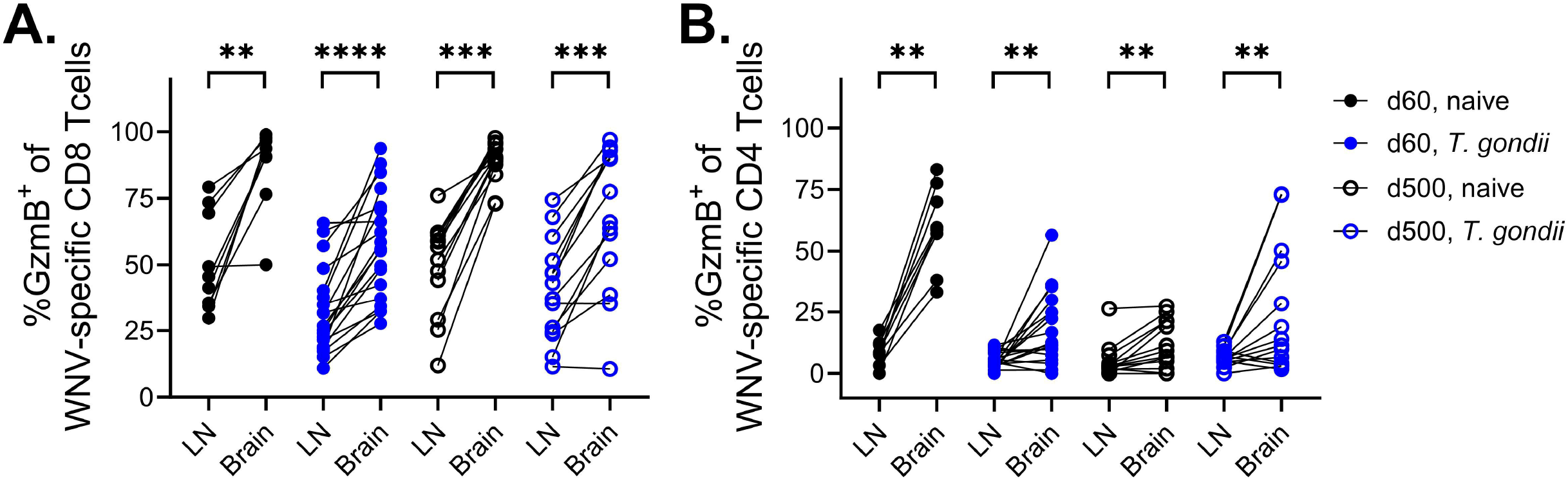

Data from both cohorts also implicate Tregs as an abundant immune cell population present in higher numbers after *T. gondii* infection, with the potential to dampen new effector responses (**Fig. 2E**). During WNV infection, Tregs expand and are thought to limit immune pathology [22,23]. To test whether resident Tregs present within the brain after *T. gondii* infection are directly limiting effector function of WNV-specific T cells, we utilized DEREG mice,where Foxp3+ Tregs express eGFP and diphtheria toxin (DT) receptors, allowing transient depletion of Tregs via DT administration. We infected DEREG mice, and their WT littermates, with *T. gondii* for 60 days, followed by WNV for 10 days as in **Fig. 1**. On days 8 and 9 post-WNV, as T cells are just arriving in the brain, we treated all mice with DT to selectively deplete Tregs and test whether we could relieve the brake on effector T cell function (schematic, **Fig. 4A**). We determined that both groups were effectively depleted of Tregs in tissue (**Fig. 4B**). We then evaluated whether depletion had an impact on WNV-specific CD8 T cell expression of GzmB. In the *T. gondii* naïve cohort, we measured that nearly 100% of all WNV-specific CD8 T cells were positive for GzmB after DT treatment, confirming the mechanistic link between Tregs and this effector function as proposed by others [22,23]. In the cohort persistently infected with*T. gondii*, we were not able to significantly improve the proportion of GzmB^+^ WNV-specific CD8 T cells **(Fig.4B**, *p*=0.0774). However, there was a clear trend towards improvement suggesting that Treg abundance in the brain may be a contributing mechanism, but perhaps not the only one, for the reduced GzmB expression by T cells in mice chronically infected with *T. gondii*.

**Figure 4.**
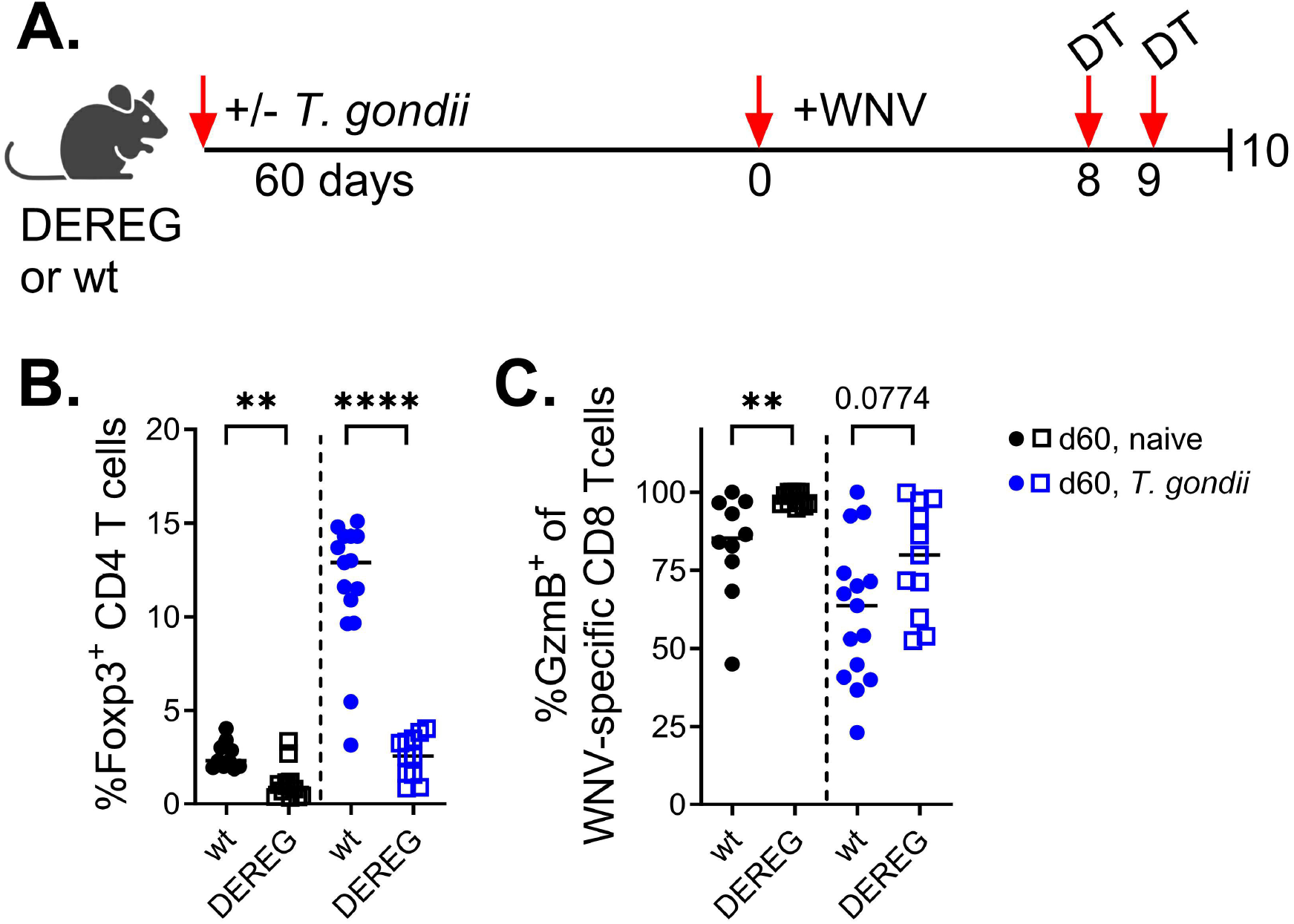

## Discussion

*T. gondii* persistence at different anatomical sites around the body, including the CNS, requires an active immune response to maintain the parasite in a quiescent state altering the local tissue milieu. We hypothesized that the presence of the parasite, and the dynamic local immune response required to maintain its encysted state, could impact acute responses to new infections with unrelated pathogens, potentially increasing host vulnerability to other neurotropic infections. Supporting this hypothesis, we found a clear association between *T. gondii* seropositivity and severe WNV disease in a clinical cohort of individuals who had been infected with WNV (**Table I**). In this cohort, those who experienced WNV encephalitis had a *T. gondii* seropositivity rate of 51% as compared to the fever-only cohort which had a rate of 19%. Both of these are considerably higher than the 11% of the U.S. population estimated by the CDC to be seropositive for *T. gondii* [4]. The increased incidence of *T. gondii* infection associated with severe WNV disease, measurable in this limited set of samples, should be further investigated in larger cohorts.

In a co-infection mouse model, we linked increased susceptibility to WNV in *T. gondii* infected mice to poor T cell effector function in the brain. This phenotype was consistent regardless of whether *T. gondii* infection was established 60 or 500 days prior demonstrating the lasting impact of parasite infection on *de novo* infection with other neurotropic pathogens.

We reveal that WNV-specific T cell effectors in the brain are poorly armed with GzmB as compared to either *T. gondii*-naïve controls or T cells in the periphery of those same mice. This finding suggested to us that a local interaction or factor was responsible for blunting GzmB activity, and this occurred even when the parasite was cleared (in 64% of mice with no detectable *T*.*gondii* at 500 days p.i.). Given the significant population of Tregs present in the brain after *T. gondii* infection, we tested our 60 day co-infection model in DEREG mice and measured a partial improvement in function when Tregs are depleted.

IFNγ is required to control tachyzoite proliferation during acute *T. gondii* infection and reactivation from encystment in the brain [24,25]. Several studies also suggest a role for granule-mediated cytotoxicity in clearing encysted parasites via T cell and perforin dependent mechanisms [10,26]. Therefore, direct, and local, inhibition of cytolytic activity would provide an obvious benefit to the parasite. *In vitro* studies have shown that *T. gondii* releases GzmB inhibitors from the parasitophorous vacuole to protect host cells from granzyme-induced apoptosis via non-serine protease mediated mechanisms [11]. In mice exclusively infected with *T. gondii*, the presence of an inhibitory factor within the tissue that limits cytotoxicity would not be readily apparent. In fact, a low amount of GzmB measured in *T. gondii*-specific cytotoxic T cells could easily be interpreted as a defect in the T cells themselves. In our study, decreased GzmB expression in WNV-specific T cells provides evidence that differentiation of antiviral T cells is specifically blunted in brain infected with *T. gondii* as compared to peripheral responses in the same mice or to the responses in *T. gondii*-naïve mice. Taken together, these data suggest that the production of local inhibitors to the granule-exocytosis pathway by cells infected with *T. gondii* may be an unidentified immune evasion mechanism.

Prevalence of *T. gondii*, and other persistent pathogens, increases with age as these infections are often acquired over the lifespan. Older adults have greater susceptibility to many acute and previously unencountered infections including WNV, SARS-CoV-2, and Influenza.

Indeed, our cohort that experienced WNV encephalitis was older than the cohort diagnosed with milder disease. The mechanisms contributing to increased susceptibility to infection with age have generally been focused on decreased magnitudes of responses or lower functionality of immune cell populations measured in blood and secondary lymphoid sites. Our results suggest that tissue environment variation by persistent pathogens acquired during life may further negatively mediate tissue-specific acute responses to WNV. Further, this tissue environment modulation could also present a barrier to adoptive T cell therapies and/or may explain the association between *T. gondii* and increased incidence of brain cancer [27,28], points that would require further investigation.

## Methods

### Human samples

Study approval was provided by the Institutional Review Boards at the University of Arizona (Tucson, AZ; # 080000673; currently #2102460536), the Oregon Health and Science University (Portland, OR; # IRB00003007); and the University of Texas Health Science Center at Houston, TX. Exclusion criteria included known immunosuppressive pathology, stroke, cancer, or use of steroids within the last 5 years. WNV-infected donors were enrolled by the University of Texas at Houston, 2006–2009. Samples, demographics (Table S1), and symptom data were collected and analyzed as previously described [29] after the subjects provided an informed consent approved by the Committee for the Protection of Human Subjects at the University of Texas Health Science Center at Houston (HSC-SPH-03-039). Blood was drawn into heparinized Vacutainer CPT tubes (BD Bioscience) and processed fresh at respective sites per manufacturer’s recommendations. Plasma was stored at -80°C.

### Pathogens

*T. gondii* parasites were maintained in culture and prepared for intraperitoneal (i.p.) inoculation exactly as previously described [18]. Live Type II (Pru-TD Tomato-OVA) or Type III (CEP) *T. gondii* tachyzoites were counted and diluted to a concentration of 50,000 tachys/mL or 10,000 parasites in a 200uL dose. Uninfected mice received USP Grade sterile PBS. Type II *T. gondii* was used in Figures 2 and 5; Type II and III were used in Figures 3 and 4 for consistency with the available lifelong cohort. Survival was monitored as shown or for 528 days (Supp. Figure 1). qPCR for parasite burden was as in [18].

West Nile virus NY 385–99, isolated from the liver of a snowy owl, was a kind gift from Dr. Robert Tesh (University of Texas Medical Branch at Galveston). Virus was prepared as previously described [19]. Mice were infected by footpad (f.p.) route with 1000 pfu WNV/50 µl/mouse and monitored for survival up to 45 days p.i.

### Mice

Mouse experiments were carried out in strict accordance with the recommendations in the Guide for the Care and Use of Laboratory Animals of the National Institutes of Health. Protocols were approved by the Institutional Animal Care and Use Committee at the University of Arizona (IACUC 19-580, PHS Assurance No. A-3248-01. Euthanasia was by overdose of isoflurane followed by perfusion with 1X PBS. C57BL/6 background mice were used in all experiments. DEREG, C57BL/6-Tg(Foxp3-DTR/EGFP)23.2Spar/Mmjax (JAX strain # 032050), were bred in house for sufficient numbers and screened by FCM for GFP+ CD4 T cells. 1ug of diphtheria toxin (DT) was administered i.p. to both DEREG and WT littermates in Figure 5 on days 8 and 9 post-WNV infection.

### ELISA

To determine anti-Toxoplasma gondii IgG in human plasma (heparin) samples we used two commercial ELISA kits (Abcam cat# ab108776 and MyBiosource cat# MBS494548) following manufacturer’s instructions.

### Flow cytometry

LN’s and Brain were harvested after perfusion with 1X PBS. LN’s were disassociated over a 40uM filter to achieve a single cell suspension. Brain was disassociated over a 40uM filter washed thoroughly with 20mL of RPMI containing 10% FBS and an aliquot was saved for WNV titer by routine plaque assay as previously described [30]. Briefly, brain mononuclear cells were isolated by pellet through 30% Percoll (Millipore Sigma) and the single-cell suspension was stained and analyzed by flow cytometric analysis on a BD Fortessa LSR cytometer.

Staining panel included a saturating concentration of the following antibodies from BioLegend (clone): Zombie dye for viability, CD45 (30-F11), CD11b (M1/70), CD3 (17A2), CD4 (RM4-4), CD8a(53-6.7), Foxp3 (MF-14), and CD103 (2E7); and from ThermoFisher: Granzyme B (GB11).Db-NS4b tetramer was provided by the NIH Tetramer Core Facility (contract number 75N93020D00005).IAb-E641 tetramer was provided by Dr. Michael S. Kuhns. Fixation and permeabilization was done with the Foxp3/Transcription Factor Staining Buffer Set (ThermoFisher). Representative gating is shown in **Supp. Figure 2**.

## Abbreviations

CNS: central nervous system
GzmB: granzyme B
pfu: plaque-forming units
T. gondii: Toxoplasma gondii
Treg: regulatory T cells
WNV: West Nile virus

## References

1. Montoya JG, Liesenfeld O. Toxoplasmosis. Lancet. 2004;363: 1965–1976.

2. Pappas G, Roussos N, Falagas ME. Toxoplasmosis snapshots: global status of Toxoplasma gondii seroprevalence and implications for pregnancy and congenital toxoplasmosis. Int J Parasitol. 2009;39: 1385–1394.

3. Calero-Bernal R, Gennari SM, Cano S, Salas-Fajardo MY, Ríos A Álvarez-García G, et al. Anti-Toxoplasma gondii Antibodies in European Residents: A Systematic Review and Meta-Analysis of Studies Published between 2000 and 2020. Pathogens. 2023;12. doi:10.3390/pathogens12121430

4. CDC - Toxoplasmosis. 15 Jan 2021 [cited 11 May 2023]. Available: https://www.cdc.gov/parasites/toxoplasmosis/index.html

5. Suzuki Y, Orellana MA, Schreiber RD, Remington JS. Interferon-gamma: the major mediator of resistance against Toxoplasma gondii. Science. 1988;240: 516–518.

6. Yap GS, Sher A. Effector cells of both nonhemopoietic and hemopoietic origin are required for interferon (IFN)-gamma- and tumor necrosis factor (TNF)-alpha-dependent host resistance to the intracellular pathogen, Toxoplasma gondii. J Exp Med. 1999;189: 1083–1092.

7. Nishiyama S, Pradipta A, Ma JS, Sasai M, Yamamoto M. T cell-derived interferon-γ is required for host defense to Toxoplasma gondii. Parasitol Int. 2020;75: 102049.

8. Gazzinelli R, Xu Y, Hieny S, Cheever A, Sher A. Simultaneous depletion of CD4+ and CD8+ T lymphocytes is required to reactivate chronic infection with Toxoplasma gondii. J Immunol. 1992;149: 175–180.

9. Denkers EY, Yap G, Scharton-Kersten T, Charest H, Butcher BA, Caspar P, et al. Perforin-mediated cytolysis plays a limited role in host resistance to Toxoplasma gondii. J Immunol. 1997;159: 1903–1908.

10. Suzuki Y, Wang X, Jortner BS, Payne L, Ni Y, Michie SA, et al. Removal of Toxoplasma gondii cysts from the brain by perforin-mediated activity of CD8+ T cells. Am J Pathol. 2010;176: 1607–1613.

11. Yamada T, Tomita T, Weiss LM, Orlofsky A. Toxoplasma gondii inhibits granzyme B-mediated apoptosis by the inhibition of granzyme B function in host cells. Int J Parasitol. 2011;41: 595–607.

12. Pierson TC, Diamond MS. The continued threat of emerging flaviviruses. Nat Microbiol. 2020;5: 796–812.

13. Bigham AW, Buckingham KJ, Husain S, Emond MJ, Bofferding KM, Gildersleeve H, et al. Host genetic risk factors for West Nile virus infection and disease progression. PLoS One. 2011;6: e24745.

14. Shrestha B, Samuel MA, Diamond MS. CD8+ T cells require perforin to clear West Nile virus from infected neurons. J Virol. 2006;80: 119–129.

15. Shrestha B, Diamond MS. Fas ligand interactions contribute to CD8+ T-cell-mediated control of West Nile virus infection in the central nervous system. J Virol. 2007;81: 11749–11757.

16. Kitaura K, Fujii Y, Hayasaka D, Matsutani T, Shirai K, Nagata N, et al. High clonality of virus-specific T lymphocytes defined by TCR usage in the brains of mice infected with West Nile virus. J Immunol. 2011;187: 3919–3930.

17. Brien JD, Uhrlaub JL, Nikolich-Zugich J. West Nile virus-specific CD4 T cells exhibit direct antiviral cytokine secretion and cytotoxicity and are sufficient for antiviral protection. J Immunol. 2008;181: 8568–8575.

18. McGovern KE, Cabral CM, Morrison HW, Koshy AA. Aging with Toxoplasma gondii results in pathogen clearance, resolution of inflammation, and minimal consequences to learning and memory. Sci Rep. 2020;10: 7979.

19. Brien JD, Uhrlaub JL, Hirsch A, Wiley CA, Nikolich-Zugich J. Key role of T cell defects in age-related vulnerability to West Nile virus. J Exp Med. 2009;206: 2735–2745.

20. Suzuki Y, Conley FK, Remington JS. Importance of endogenous IFN-gamma for prevention of toxoplasmic encephalitis in mice. J Immunol. 1989;143: 2045–2050.

21. Denkers EY, Gazzinelli RT. Regulation and function of T-cell-mediated immunity during Toxoplasma gondii infection. Clin Microbiol Rev. 1998;11: 569–588.

22. Lanteri MC, O’Brien KM, Purtha WE, Cameron MJ, Lund JM, Owen RE, et al. Tregs control the development of symptomatic West Nile virus infection in humans and mice. J Clin Invest. 2009;119: 3266–3277.

23. Graham JB, Swarts JL, Edwards KR, Voss KM, Green R, Jeng S, et al. Correlation of Regulatory T Cell Numbers with Disease Tolerance upon Virus Infection. Immunohorizons. 2021;5: 157–169.

24. Suzuki Y, Sa Q, Gehman M, Ochiai E. Interferon-gamma- and perforin-mediated immune responses for resistance against Toxoplasma gondii in the brain. Expert Rev Mol Med. 2011;13: e31.

25. Munoz M, Liesenfeld O, Heimesaat MM. Immunology of Toxoplasma gondii. Immunol Rev. 2011;240: 269–285.

26. Tiwari A, Hannah R, Lutshumba J, Ochiai E, Weiss LM, Suzuki Y. Penetration of CD8+ Cytotoxic T Cells into Large Target, Tissue Cysts of Toxoplasma gondii, Leads to Its Elimination. Am J Pathol. 2019;189: 1594–1607.

27. Vittecoq M, Elguero E, Lafferty KD, Roche B, Brodeur J, Gauthier-Clerc M, et al. Brain cancer mortality rates increase with Toxoplasma gondii seroprevalence in France. Infect Genet Evol. 2012;12: 496–498.

28. Abdollahi A, Razavian I, Razavian E, Ghodsian S, Almukhtar M, Marhoommirzabak E, et al. Toxoplasma gondii infection/exposure and the risk of brain tumors: A systematic review and meta-analysis. Cancer Epidemiol. 2022;77: 102119.

29. Lanteri MC, Lee TH, Wen L, Kaidarova Z, Bravo MD, Kiely NE, et al. West Nile virus nucleic acid persistence in whole blood months after clearance in plasma: implication for transfusion and transplantation safety. Transfusion . 2014;54: 3232–3241.

30. Uhrlaub JL, Brien JD, Widman DG, Mason PW, Nikolich-Zugich J. Repeated in vivo stimulation of T and B cell responses in old mice generates protective immunity against lethal West Nile virus encephalitis. J Immunol. 2011;186: 3882–3891.

