## Supplemental Figures 1 and 2 for "Chronic *Toxoplasma gondii* infection curtails the cytotoxic potential of acute T cell responses to West Nile virus in the brain"

**Supplemental Figure 1**

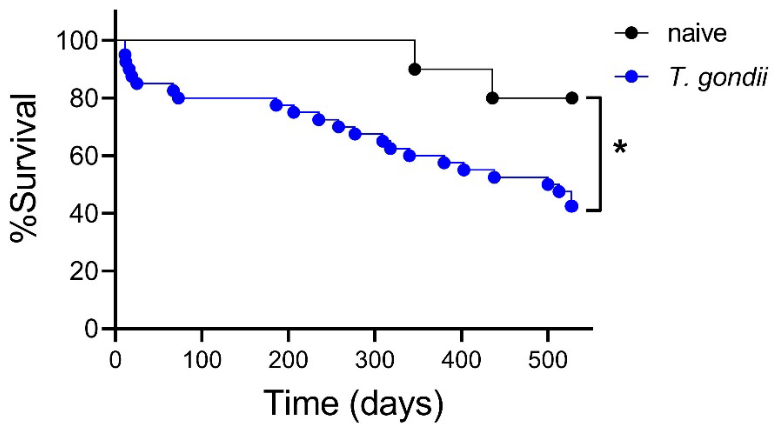


***T. gondii* infection negatively impacts survival over lifespan.**

C57BL/6 mice (mixed male and female, 3mo) were challenged with *T. gondii* (10,000 tachyzoites, ip) to establish *T. gondii* infection alongside relevant naïve controls (saline, ip) and monitored for survival. *T. gondii* (n=40) vs. naïve controls (n=10) at day 60 post-*T. gondii* infection P=0.0400,*. Significance of survival evaluated by Log-rank (Mantel-Cox) test. Splitting the data by gender reduces power and statistical significance between groups is lost. Median survival of the *T. gondii* infected cohort was 506.5 days.

**Supp. Figure 2**

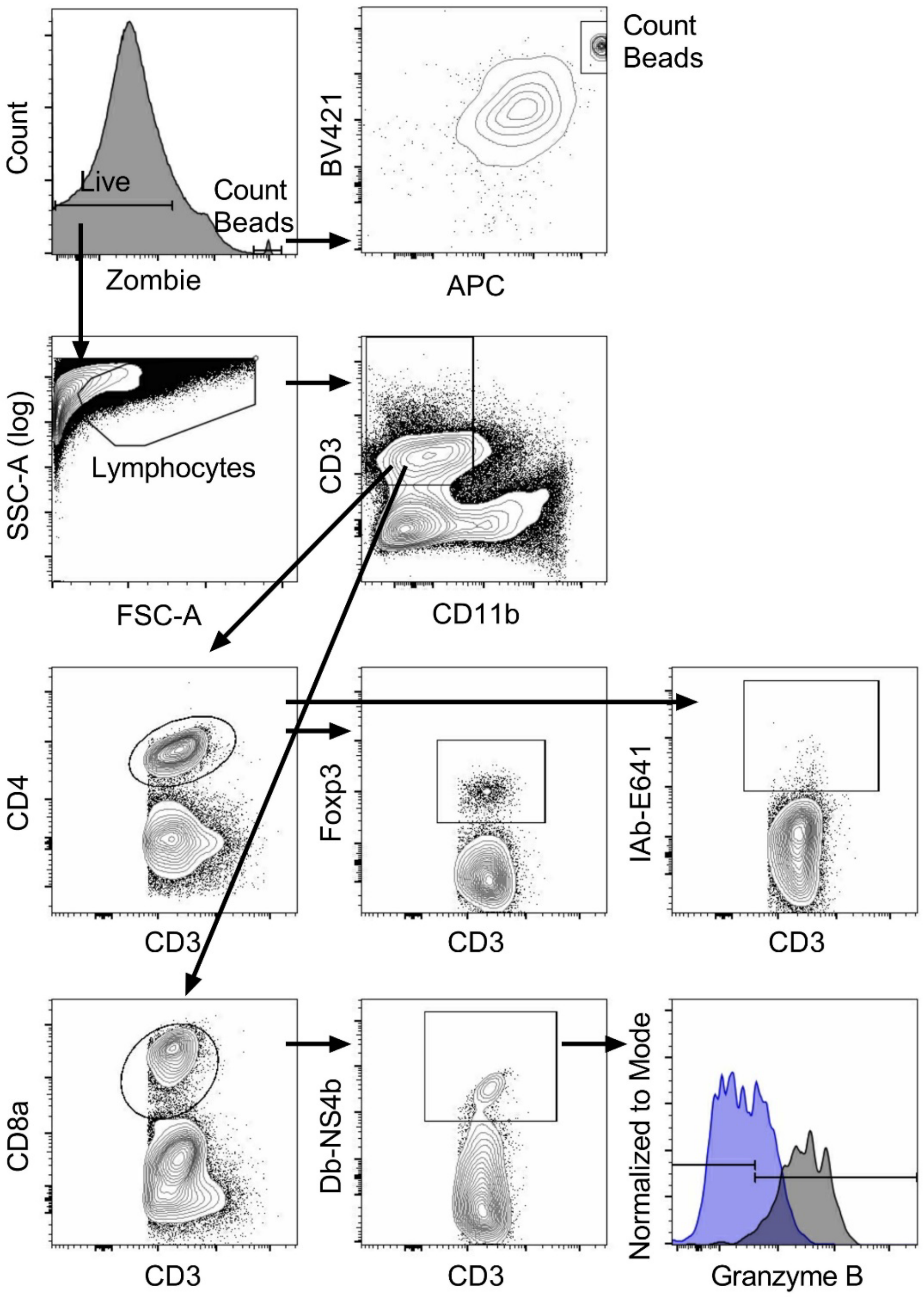


**Representative Flow Cytometry Gating**

Sample shown is brain from a lifelong *T. gondii* infected mouse on day 10 post-WNV. For the bottom right panel, WNV-specific CD8 T cells are gated as positive or negative based on histogram for Granzyme B; dark blue is the d500 *T. gondii* infected mouse consistent with the rest of the layout overlayed by a d500 *T. gondii naïve* mouse on day 10 WNV to show the change in Granzyme B expression. Data shown is from Figure 2.
